# Innervated adrenomedullary microphysiological system to model prenatal nicotine and opioid exposure

**DOI:** 10.1101/2020.09.22.308973

**Authors:** Jonathan R. Soucy, Gabriel Burchett, Ryan Brady, David T. Breault, Abigail N. Koppes, Ryan A. Koppes

## Abstract

The transition to extrauterine life results in a critical surge of catecholamines necessary for increased cardiovascular, respiratory, and metabolic activity. The mechanisms mediating adrenomedullary catecholamine release are poorly understood, given the sympathetic adrenomedullary control systems’ functional immaturity. Important mechanistic insight is provided by newborns delivered by cesarean section or subjected to prenatal nicotine or opioid exposure, demonstrating the impaired release of adrenomedullary catecholamines. To investigate mechanisms regulating adrenomedullary innervation, we developed compartmentalized 3D microphysiological systems (MPS) by exploiting the meniscus pinning effect via GelPins, capillary pressure barriers between cell-laden hydrogels. The MPS comprises discrete 3D cultures of adrenal chromaffin cells and preganglionic sympathetic neurons within a contiguous bioengineered microtissue. Using this model, we demonstrate that adrenal chromaffin innervation plays a critical role in hypoxia-medicated catecholamine release. Furthermore, opioids and nicotine were shown to affect adrenal chromaffin cell response to a reduced oxygen environment, but neurogenic control mechanisms remained intact. GelPin containing MPS represent an inexpensive and highly adaptable approach to study innervated organ systems and improve drug screening platforms by providing innervated microenvironments.

## INTRODUCTION

Catecholamines (CATs) synthesized in the adrenal medulla regulate tissue and organ homeostasis. In response to physical or psychological stress, the adrenal medulla rapidly releases high concentrations of CATs, increasing blood plasma levels (>300-fold increase), commonly known as the ‘fight or flight’ response (1, 2). Anatomically, cholinergic preganglionic nerve fibers of the sympathetic distribution innervate the adrenal medulla and synapse with adrenomedullary chromaffin cells (AMCCs) to facilitate CAT release (3). These preganglionic sympathetic neurons (PSNs) reside in the medial horn of the spinal cord (T5-T8) and mediate the dynamic response to external and internal stimuli (4). Innervation of the adrenal medulla, however, does not mature until early postnatal life. During a vaginal delivery, in response to hypoxia and hypercapnia, an AMCC CAT surge facilitates the transition to extrauterine life (5, 6). These nonneurogenic mechanisms regress postnatally (7).

During the perinatal transition, CATs enter the bloodstream via the inferior vena cava, are distributed systemically, and bind to adrenergic receptors expressed in the heart and lungs to promote cardiovascular adaptation (8, 9). CATs approach their highest concentrations in life during vaginal delivery, but CAT concentrations are significantly reduced for newborns delivered by cesarean section (10). With the rise of cesarean sections in recent decades, this attenuated CAT surge has been proposed as a potential mechanism leading to perinatal cardiorespiratory concerns in this population (11–15). Prenatal exposure to nicotine and opioids also impairs the perinatal CAT surge in newborns independent of the route of delivery (16, 17). However, contemporary model systems cannot elucidate the underlying cellular mechanisms resulting from such exposures. The regulation of CAT levels is also necessary during postnatal life for increasing the cardiovascular, respiratory, and metabolic activity in response to acute stress (e.g., the ‘flight-or-fight’ response) (3, 10, 18); however, the long-term ramifications of a depressed response at birth remains unclear. Further, prenatal exposure to drugs of abuse may impact both short- and long-term neurohormonal control systems. Specifically, low CAT levels at birth can significantly delay or impede key development milestones (19).

In addition to the impact of adrenomedullary dysfunction at birth, a strong correlation exists between high levels of emotional and physical stress and cardiovascular disease (CVD) (20). The ‘flight-or-fight’ response originally evolved as a survival trait to handle acute physical stress but has become maladaptive in light of the chronic psychological stressors characteristic of modern human society (21). Furthermore, continuous activation of the adrenal gland results in poor cardiovascular health (22) and has been associated with chronic fatigue syndrome, depression, and post-traumatic stress disorder (23). Therefore, new tools to investigate adrenal biology are critical for understanding human health and disease.

To date, the studies of CAT release, using either AMCCs or adrenal slice culture, often ignore the role of innervation from the sympathetic adrenomedullary (SAM) axis (24, 25). Once isolated from animals or differentiated from stem cells, both AMCCs and cultured adrenal slices lack neural regulation. Both *in vivo* and *ex vivo* models have been used to investigate the SAM axis, and together have helped establish our current understanding of neurogenic control of CAT release (26). Due to the confounding impact of autonomic regulation and other feedback mechanisms, however, *in vivo* approaches are ill-suited for developing a comprehensive understanding of the cellular mechanisms regulating CAT release. In contrast, *in vitro* models, with their ability to reduce complexity, control for variability, and increase experimental throughput, may be used in place of animal models for discovery applications (27).

Microfluidic devices provide an attractive opportunity for the development of innervated organ systems *in vitro* because of their capacity to compartmentalize cells and incorporate real-time measures of cell function (28, 29). To date, these tools have not been utilized to model the SAM axis. Primary bovine AMCCs have been cultured in a 2D microfluidic platform for measuring CAT release in response to secretagogues (30); however, this model failed to mimic the 3D physiological microenvironment of the adrenal medulla and lacked neural inputs. Cellular morphology and phenotype are strongly influenced by the chemical, mechanical, and cell-cell interactions within the 3D multicellular microenvironment of endogenous tissues (31); characteristics that 2D cell culture systems often fail to capture. Moreover, previous microfluidic systems used to assess CAT release were most commonly fabricated using polydimethylsiloxane (PDMS), which is gas permeable and known to absorb small and charged molecules (32–35). Therefore, to better recapitulate the *in vivo* environment of the SAM axis, we engineered a thermoplastic microphysiological system (MPS) to co-culture AMCCs and PSNs in a biomimetic scaffold.

Here, we utilize an adaption of our previously described ‘cut & assemble’ process for manufacturing thermoplastic MPS to investigate mechanisms regulating CAT release by innervated AMCCs (36, 37). Furthermore, our MPS designs feature gas impermeability, allowing control over oxygen tension needed to mimic the hypoxic environment present during the perinatal transition (32–35). Traditionally, compartmentalization is established using either micro-posts or micro-tunnels to constrain specific cell populations to discrete regions (38–40). To facilitate similar spatial control without the need for high-resolution manufacturing, we exploited the meniscus pinning effect with thin floating guides, dubbed GelPins, to establish temporary liquid-air interfaces within our device via photocrosslinkable hydrogels (Fig. 1) (37). By sequentially loading the MPS with hydrogel precursors containing either AMCCs or PSNs, each solution can be crosslinked in situ to form a contiguous gel layer with discrete cellular compartmentalization within a multilayer microfluidic device. Sympathetic innervation was validated by neurite growth towards and synapsis with AMCCs, and CAT release at physiological concentrations was assessed using cardiomyocytes (CMs). Altogether, our adrenomedullary MPS represents a versatile platform to evaluate the impact of acute stressors on CAT release via establishing a neuro-adrenal unit *in vitro*. Furthermore, a detailed understanding of the mechanisms regulating the SAM axis and AMCCs may lead to the elucidation of novel therapeutic targets for the treatment of CVD and other stress-related disorders.

**Fig. 1.**
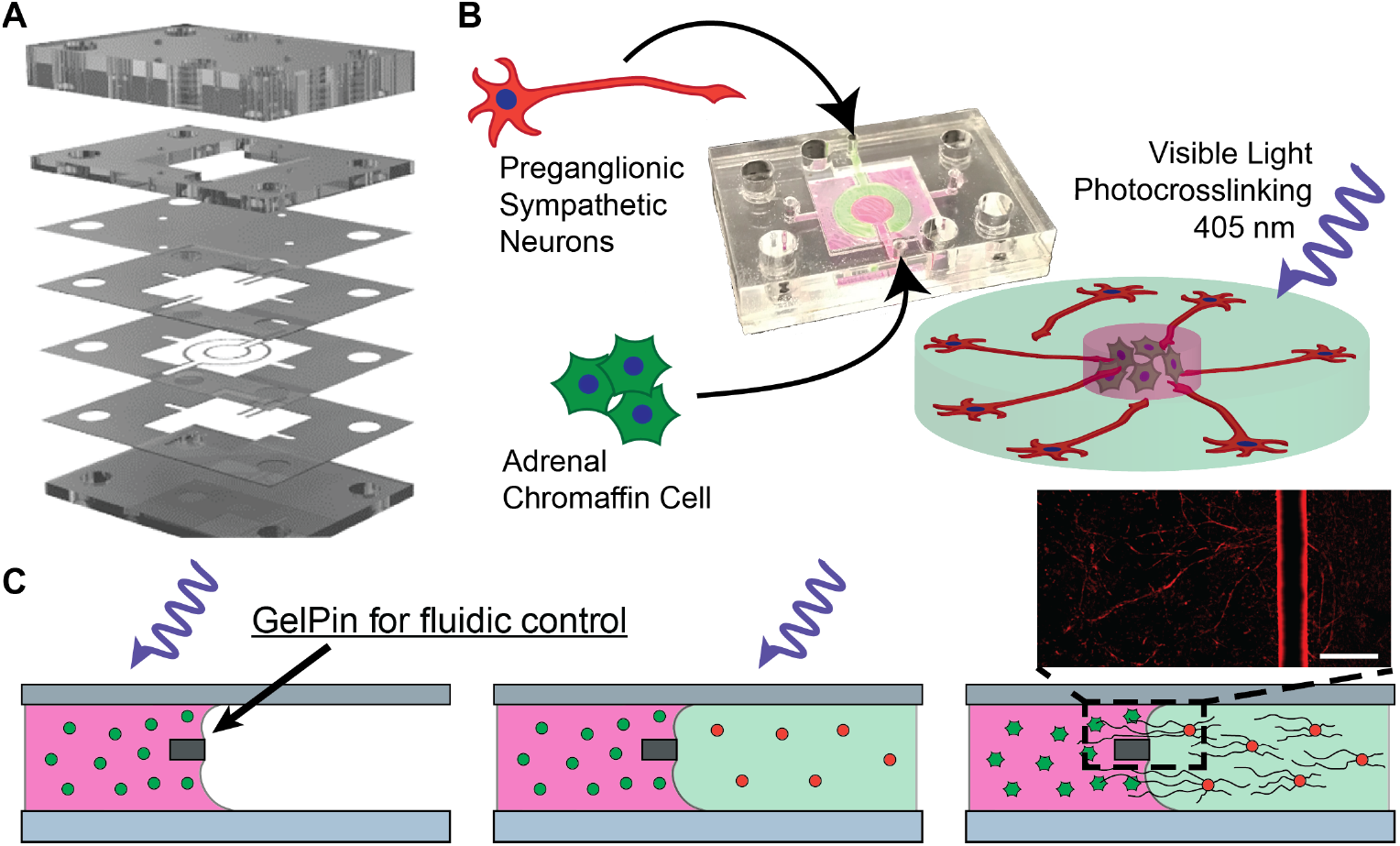
Adrenal MPS schematic. (**A-B**) Multilayer chip assembly for the 3D culture of preganglionic sympathetic neurons and adrenal chromaffin cells within a photocrosslinkable hydrogel. (**C**) Cross-sectional representations of step-by-step concentric-circle MPS (ccMPS) assembly resulting in a contiguous hydrogel with discrete compartmentalization. Representative immunofluorescent image showing neurites (βIII tubulin, red) crossing the GelPin (autofluorescence). Scale = 300 μm.

## RESULTS

### Development of an innervated adrenomedullary MPS

To expand the utility of our ‘cut & assemble’ manufacturing technique to model multi-tissue interactions (36), we established GelPins to control cell-laden microtissues’ size and shape via the meniscus pinning effect within our devices (Fig. 1A). We generated an innervated adrenomedullary MPS consisting of the following seven layers irreversibly bonded together: (1) a top 4.5 mm layer that seals the MPS and provides fluidic connections, (2) a 1.5 mm apical flow channel for circulation of media, (3) a CycloporeTM transparent polycarbonate membrane (1 *μ*m pore size) to interface the basal and apical channels and allow for nutrient diffusion into the microtissue, (4) a 60 *μ*m contiguous gel compartment to enable discrete gel contacts, (5) a 76 *μ*m GelPin to enable the formation of individual gels with defined geometries, (6) a 120 *μ*m contiguous gel compartment providing an additional gel interfacing area, and (7) a glass coverslip to seal the bottom of the MPS and enable high-resolution imaging. Each layer’s geometry was designed in computer-aided drafting (CAD) software and cut out using a commercially available CO_2_ laser engraver.

The use of floating GelPins allows multiple discrete hydrogels to be seeded adjacent to one another, forming a contiguous hydrogel layer with a predetermined thickness (~250 *μ*m) in the basal layer of the SAM bi-layer MPS (Fig. 1B). Unlike microfluidic devices using micro-posts or micro-tunnels to establish compartmentalization, our custom platforms enclose unobstructed, well-defined interfaces between compartments that can better mimic native tissue interfaces. For instance, the concentric-circle MPS (ccMPS) design contains regions to culture cells from both the adrenal medulla and neurons from the SNS (Fig. 1C). Immunostaining of the MPS demonstrates viable encapsulated cells and neurites extending across the floating GelPin towards the AMCCs while maintaining physiological compartmentalization of cell bodies (Fig. 1C, inlay). Further, the individual gel compartments may be filled on separate days to allow for additional preparation time or maturation of individual cell populations. Lastly, the membrane separating the basal and apical channels of the bi-layer chip design enables homogenous, circulation-like diffusion of nutrients without generating the chemotactic gradient typical of devices with flanking media channels (41, 42).

### Characterization of PNS and AMCC co-culture

*In vivo*, cholinergic preganglionic nerve fibers, originating in the intermediolateral column of the sympathetic division of the autonomic nervous system, innervate the adrenal medulla to regulate the release of epinephrine (EP) and norepinephrine (NEP) into the systemic circulation (Fig. 2A) (4). To develop a SAM MPS, we first isolated primary neonatal rat PSNs and AMCC. We then confirmed their phenotypes using immunofluorescent staining for chromogranin A (CHGA), phenylethanolamine N-methyltransferase (PNMT), nitric oxide synthase (nNOS), calretinin, and choline acetyltransferase (ChAT) 5 days post-dissociation (Fig. 2B-C). Neonatal cells were chosen to establish a developmental model of the SAM axis because neural modulation of adrenomedullary function is absent until ~7 – 10 days of postnatal life in the rat (18). Furthermore, the use of neonatal rat cells eliminates the confounding influence of innervation *in vivo* and an endogenous response to a reduced oxygen environment (18). To confirm these AMCCs have the capacity to secrete CATs, we recorded extracellular action potentials, as previously described (30). In brief, the presence of AMCC depolarizations, a surrogate for CAT exocytosis, was monitored using a multi-electrode array (MEA) following stimulation with pituitary adenylate cyclase-activating peptide (PACAP) (Fig. 2D). For all groups, we recorded baseline spontaneous depolarization events (Fig. S1, Fig. 2D), which were increased 2 – 3-fold following PACAP stimulation. Specifically, AMCCs demonstrated a baseline of 6.8 ± 2.3 spikes/min, which increased to 17.4 ± 8.0 spikes/min 5 – 10 min following PACAP exposure (p < 0.05). Finally, AMCC depolarization was attenuated following the removal of PACAP from the media, returning to a baseline of 6.9 ± 2.0 spikes/min, demonstrating temporal control of the model system.

**Fig. 2.**
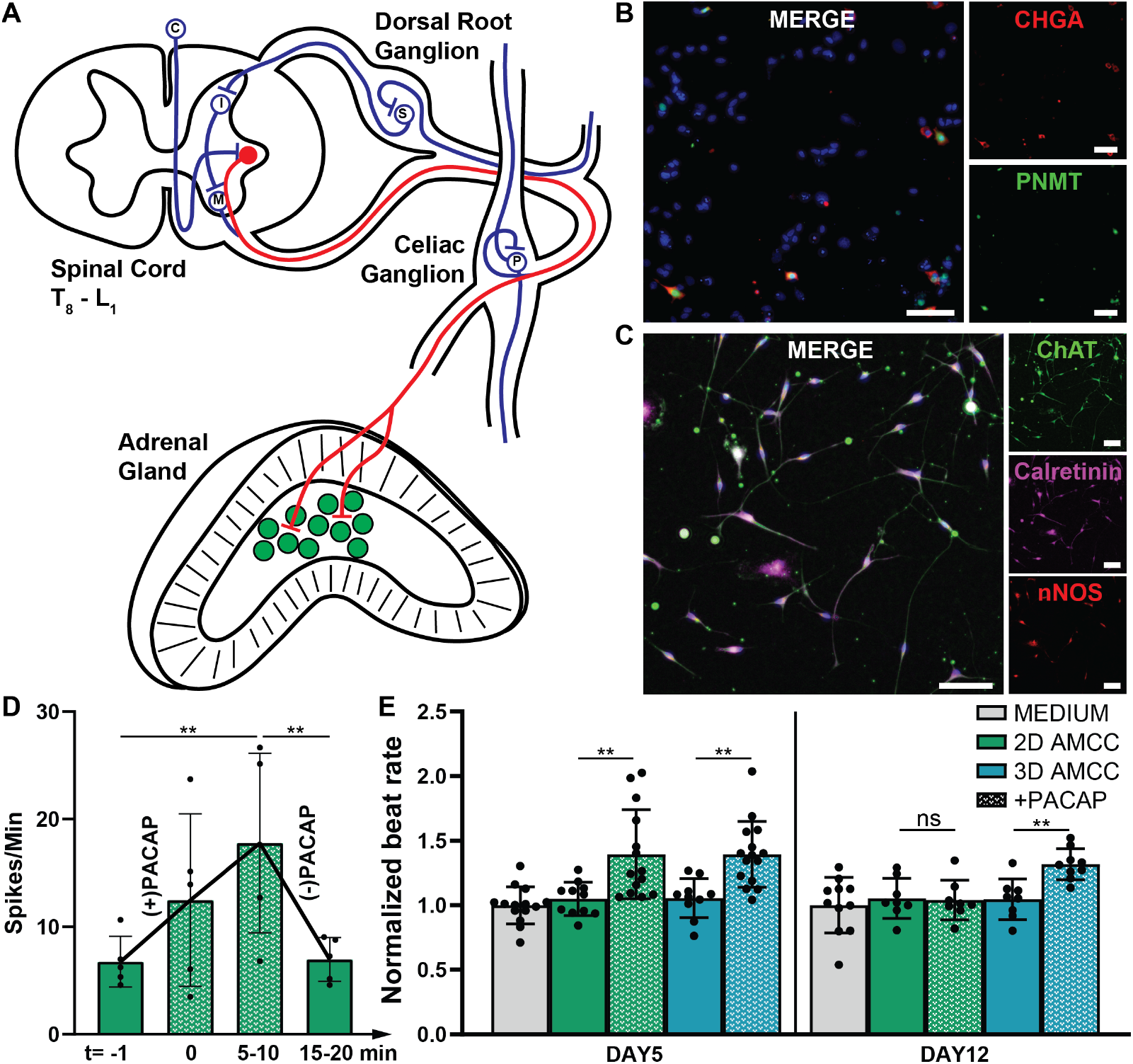
Characterization of PSN and AMCC culture. (**A**) Diagram indicating where each specific cell population was isolated. (**B**) Representative immunofluorescent images of primary AMMCs showing immunostaining for CHGA (red), PNMT (green), and DAPI (blue) and (**C**) primary PSNs showing immunostaining for ChAT (green), calretinin (violet), nNOS (red), and DAPI (blue). Scale = 100 μm. (**D**) The number of AMCC depolarization events significantly increases in response to PACAP stimulation within 5 – 10 minutes but is returned to baseline following removal of PACAP. (**E**) AMCCs cultured in 2D and 3D respond to PACAP on day 5, but only those in 3D continue to be responsive at 12 days in culture. **p < 0.05, ns: not significant

Cardiomyocytes (CMs) rapidly respond to changes in CAT levels, both *in vivo* and *in vitro*. Unlike bioelectronic sensors that oxidize CATs to calculate a concentration, CMs act as a biological transducer without altering CAT concentration. CMs quickly respond to small changes in CAT concentrations, exhibited via altered beat rate (Fig. S2). We used video microscopy and a custom cell-by-cell image analysis algorithm (43) to calculate changes in CM beat rate in response to changes in CAT concentrations. This approach represents a new experimental approach to correlate SAM innervation with catecholamine exocytosis changes *in vitro* (Fig. S3). AMCC CAT exocytosis was investigated in both 2D and 3D. AMCCs and PSNs were encapsulated in a gelatin methacrylate (GelMA) hydrogel to mimic the *in vivo* microenvironment and maintain their function during long-term culture. After five days in culture, PACAP stimuli on both 2D and 3D AMCC cultures resulted in a 39% increase in beat rate compared to controls (Fig. 2E). In contrast, after 12 days, AMCCs cultured in 2D no longer respond to PACAP stimulation, while those cultured in 3D continued to release CATs in response to PACAP and showed an increased beat rate of 32%, thereby highlighting the relevance of 3D culture systems for long-term studies.

The adrenal medulla is surrounded by the adrenal cortex and is continuously exposed to high glucocorticoid concentrations due to centripetal blood flo w. Glucocorticoids upregulate EP production by inducing expression of phenylethanolamine N-methyltransferase (PNMT) within AMCCs. To assess the effect of glucocorticoids on AMCC CAT exocytosis, culture media was supplemented with dexamethasone (DEX), a potent ligand of the glucocorticoid receptor, and both EP and NEP concentrations were quantified. Compared with untreated controls, DEX treated cultures showed an 85 ± 36% increase in EP levels in response to PACAP, while no significant change in NEP levels was observed (Fig. 3A), indicating an increase in the EP/NEP ratio. Of note, treatment with DEX did not alter the overall CM response to PACAP-induced AMCC exocytosis (Fig. 3B). Immunofluorescent images were analyzed to determine the influence of DEX on chromogranin A (CHGA+) expression (Fig. 3C), an indication of higher chromaffin granule concentration and CATs for release. In the presence of PACAP, CHGA+ expression is observable in a higher proportion of cells following DEX treatment (Fig. 3C). However, despite this immunofluorescent indication of increased CAT capacity and release, the percentage of CAT released in response to PACAP did not change. Lastly, immunostaining revealed no significant changes in the colocalization of CHGA+ and PNMT in response to DEX (Fig. S4).

**Fig. 3.**
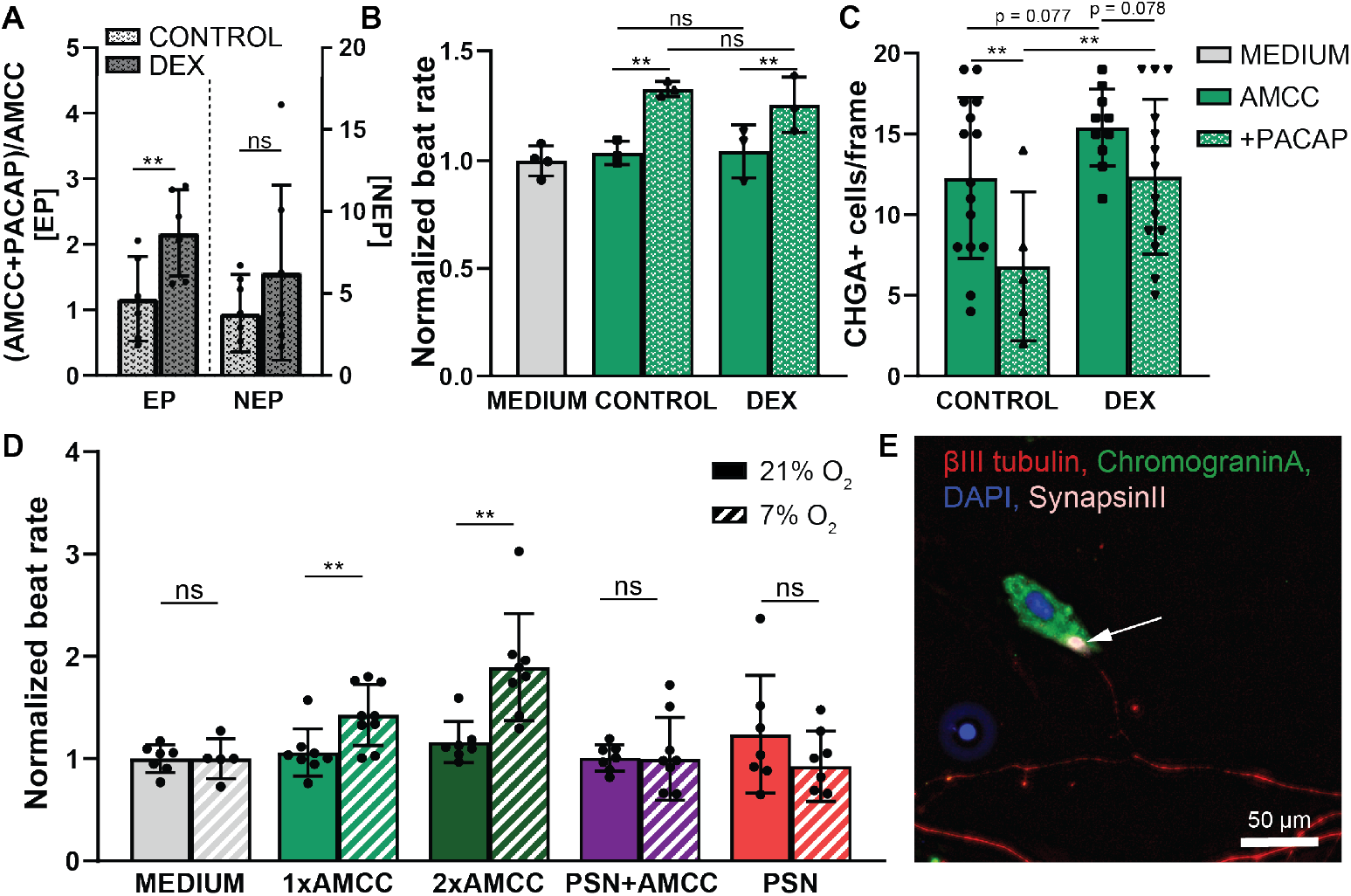
Recapitulating the adrenal microenvironment *in vitro* through pharmacological manipulation and neuronal co-culture. **(A)** DEX treatment increases the percentage AMCC EP release in response to PACAP, while not significantly altering NEP exocytosis, **(B)** but does not alter the CM response to PACAP-induced AMCC exocytosis. **(C)** DEX increases the total number of chromogranin A positive (CHGA+) cells/frame compared to controls for both PACAP stimulated (p = 0.038) and non-stimulated (p = 0.077) AMCCs. However, the addition of PACAP reduces CHGA+ cells/frame for both Controls (p = 0.044) and DEX (p = 0.078) treated AMCCs, indicating exocytosis. **(D)** CM beat rate increases in response to conditioned media from AMCCs at 500 (1x) or 1000 (2x) cells/mm^2^ exposed to rO_2_, but not in response to conditioned media from AMCC+PSN coculture (500 AMCCs/ mm^2^ + 500 PSNs/mm^2^) nor PSN monoculture (500 PSNs/mm^2^). **(E)** Representative image of PSN+AMCC co-culture showing Synapsin II (white) positive connections between an AMCC (green) and a PSN (red) after five days in culture. **p < 0.05, ns: not significant

### Modeling adrenomedullary synaptogenesis and low oxygen *in vitro*

To investigate nonneurogenic mechanisms mediating catecholamine release, we modeled the ~2 – 3-fold reduction in oxygen tension [reduced oxygen conditions, (rO_2_): 7 vol% O_2_] that occurs during the transition from intrauterine to extrauterine life (5, 44). In contrast to PDMS-based microfluidic devices, the lower gas permeability of MPS platforms allows for the effects of altered oxygen tension to be investigated without the need for specialized capital equipment or material modification (45). Conditioned media from AMCCs cultured at 500 cells/mm^2^ (1x) and exposed to rO_2_ for 4 hours showed a 43% increase in CM beat rate, while conditioned media from AMCCs cultured at 1000 cells/mm^2^ (2x) resulted in a 90% increase compared to baseline (Fig. 3D). In contrast, no changes in CM beat rate were observed using conditioned media from AMCC+PSN co-cultures exposed to rO_2_ compared to normoxia controls (Fig. 3D). Finally, immunocytochemical analysis confirms that co-culture leads to PSN neurite extensions and synaptic junctions with AMCCs (Fig. 3E). Together, these data suggest that AMCC innervation by PSNs inhibits catecholamine exocytosis in response to rO_2_, highlighting the importance of including SAM in adult MPS platforms.

### Role of nicotine and opioid exposure on adrenomedullary CAT exocytosis

Clinical evidence suggests that prenatal exposure to drugs of abuse such as nicotine and opioids can lead to low plasma levels of CATs during the transition to extrauterine life (46). How exposure to these compounds influences AMCCs or sympathetic development during gestation remains unclear. To investigate the impact of nicotine and opioid exposure, AMCCs were treated with either nicotine (1 *μ*M) or a *μ*-opioid agonist (Tyr-D-Arg-Phe-Lys, DALDA) (1 *μ*M). Compared to control cultures, exposure to either nicotine or DALDA inhibited the rO_2_-induced increase in CM beat rate following exposure to conditioned media (Fig. 4A).

**Fig. 4.**
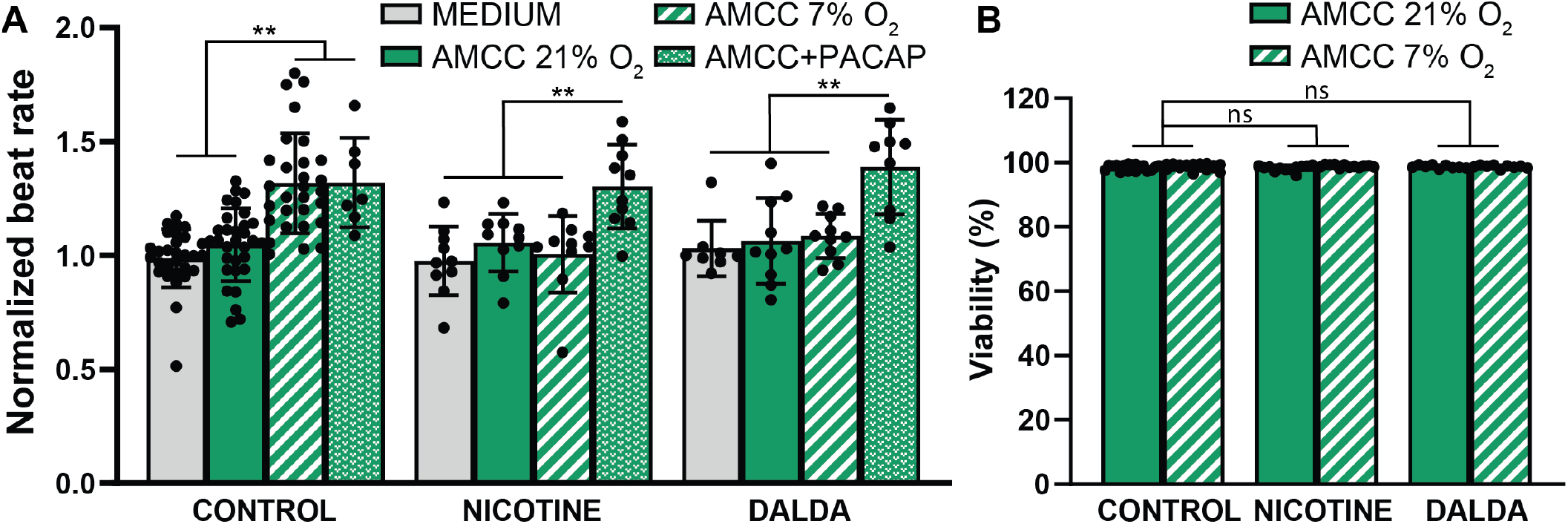
Drugs of addiction influence catecholamine exocytosis. **(A)** Conditioned media from AMCCs exposed to rO_2_ (4 hours) or PACAP results in an increased CM beat rate. In contrast, exposure to nicotine or DALDA prevents the rO_2_-induced increase in CM beat rate but does not affect PACAP responsiveness. **(B)** Nicotine, DALDA, nor rO_2_ exposure do not impact AMCC viability. **p < 0.05, ns: not significant

PACAP binds to the PAC1 receptor to trigger AMCC CAT release in response to stress (47, 48). However, unlike hypoxia-induced CAT exocytosis, PACAP-induced CAT exocytosis does not rely on membrane depolarization (49). Prolonged nicotine and opioid exposure likely inhibit CAT exocytosis in response to rO_2_ due to exposure induced alterations in AMCC ion channel expression (50, 51). Therefore, to confirm PAC1 receptor function remains unaffected by drug exposure, AMCCs CAT exocytosis was quantified in response to PACAP induction following nicotine or DALDA exposure. PACAP induction resulted in a similar concentration of CATs compared to controls, as shown by the > 30% increase in CM beat rate for each group (Fig. 4A). Thus, in addition to confirming PAC1 receptor function, drugs of abuse do not appear to diminish CAT synthesis. Finally, to confirm that the observed differences in CAT release were not a result of cell death, we studied the impact of DALDA or nicotine exposure under normoxic or rO_2_ conditions. No differences in cell viability were detected (Fig. 4B).

## DISCUSSION

To date, no 3D culture systems of the sympathetic adrenomedullary (SAM) axis have been presented. The adrenal medulla has a significant impact on plasma catecholamine levels, which influences most organ systems. Several pharmaceutical compounds designed to improve the prognoses of neurological disorders, including attention deficit hyperactivity disorder, narcolepsy, epilepsy, depression, and even obesity, have unintended side effects that result in altered CAT levels, which may require additional clinical interventions (52, 53). These unintended results are a direct result of our poor understanding of the SAM axis and the underlying mechanisms regulating AMCCs. Despite the urgent need to better understand the SAM axis, few attempts to decouple AMCC function from the complex autonomic regulation present *in vivo* have been presented. Such a model is paramount for fundamental discovery or drug screening applications. Regardless of the nervous system’s critical role for tissue development and function, innervation is often overlooked in most tissue engineering applications due to the complexities of handling multiple, highly sensitive cell populations and tissue structures with traditional processing techniques (25).

Towards establishing a reproducible system that would facilitate both 3D culture and innervation, we describe the utilization of a new, inexpensive, compartmentalized MPS, which recapitulates essential features of the sympathetic innervation of the adrenal medulla (Fig. 1). The ‘cut & assemble’ manufacturing technique provides rapid (hours), simple, and inexpensive (< $2 per device) access to multilayer 3D organs-on-chips with standard fluidic connectors (36). Notably, such an approach enables rapid and inexpensive prototyping for iterative design and increased access to organ-chip systems. Further, thermoplastic chip construction should minimize analyte loss, including CATs, compared to traditional PDMS-based devices (34, 35). The materials utilized herein allow for high optical clarity and light transmission enabling real-time, high-resolution microscopic monitoring of cellular processes (45). Finally, transparency facilitates rapid and spatiotemporal photopolymerization of hydrogels on-chip, thereby increasing reliability, experimental throughput, and achievable architectures (54, 55).

Culturing in 3D is critical for maintaining long-term functionality of adrenomedullary cells *in vitro*. AMCCs encapsulated within GelMA hydrogels sustain CAT exocytosis in response to PACAP or neurogenetic stimuli after 12 days in culture, whereas the response is lost in AMCCs cultured in 2D (Fig. 1E). Our MPS design may be used to establish oxygen gradients to better mimic the anaerobic environments of some tissues without additional processing (56). AMCCs co-cultured with PSN does not exhibit CAT exocytosis in response to reduced oxygen tension, a prenatal reflex. Our *in vitro* data suggest that synaptogenesis, or possibly just the presence of cholinergic preganglionic sympathetic neurons, is responsible for inhibiting the rO_2_-induced release of catecholamines (Fig. 3D). These results highlight the sensitivity of AMCCs to oxygen levels and the importance of recapitulating the SAM axis in an adrenomedullary MPS to more faithfully recapitulate physiological control mechanisms.

Current working models of rO_2_-induced CAT exocytosis suggest that a low oxygen environment during natural birth leads to a reduction of H_2_O_2_ as part of the mitochondria’s regulation of oxidative stress (Fig. 5) (7, 57). This reduction in H_2_O_2_ results in a closing of low and high conductance potassium (SK/BK) ion channels, which triggers a depolarization event. In response to this membrane depolarization, voltage-gated calcium (Ca^2+^) channels open. The increase in intracellular calcium then promotes CAT exocytosis. When the prenatal adrenal medulla is exposed to opioids and nicotine; however, AMCCs will overexpress adenosine triphosphate (ATP)-sensitive potassium (KATP) channels (Fig. 5) (50, 51). Moreover, these KATP channels will remain open in a low oxygen environment due to limited ATP availability, thereby preventing depolarization and subsequent hypoxia-induced CAT release. In our model, exposure to either a-opioid agonist or nicotine attenuated the response to rO_2_ but did not prevent neurogenetic control of CAT exocytosis, suggesting a potential target to ameliorate the loss of CAT surge during birth. Additionally, while nicotine appears to prevent the CAT surge caused by rO_2_ directly, nicotine also increases neurite growth rate (58), which may accelerate adrenomedullary synaptogenesis before birth and bolster the resulting loss of CAT exocytosis (Fig. 3D).

**Fig. 5.**
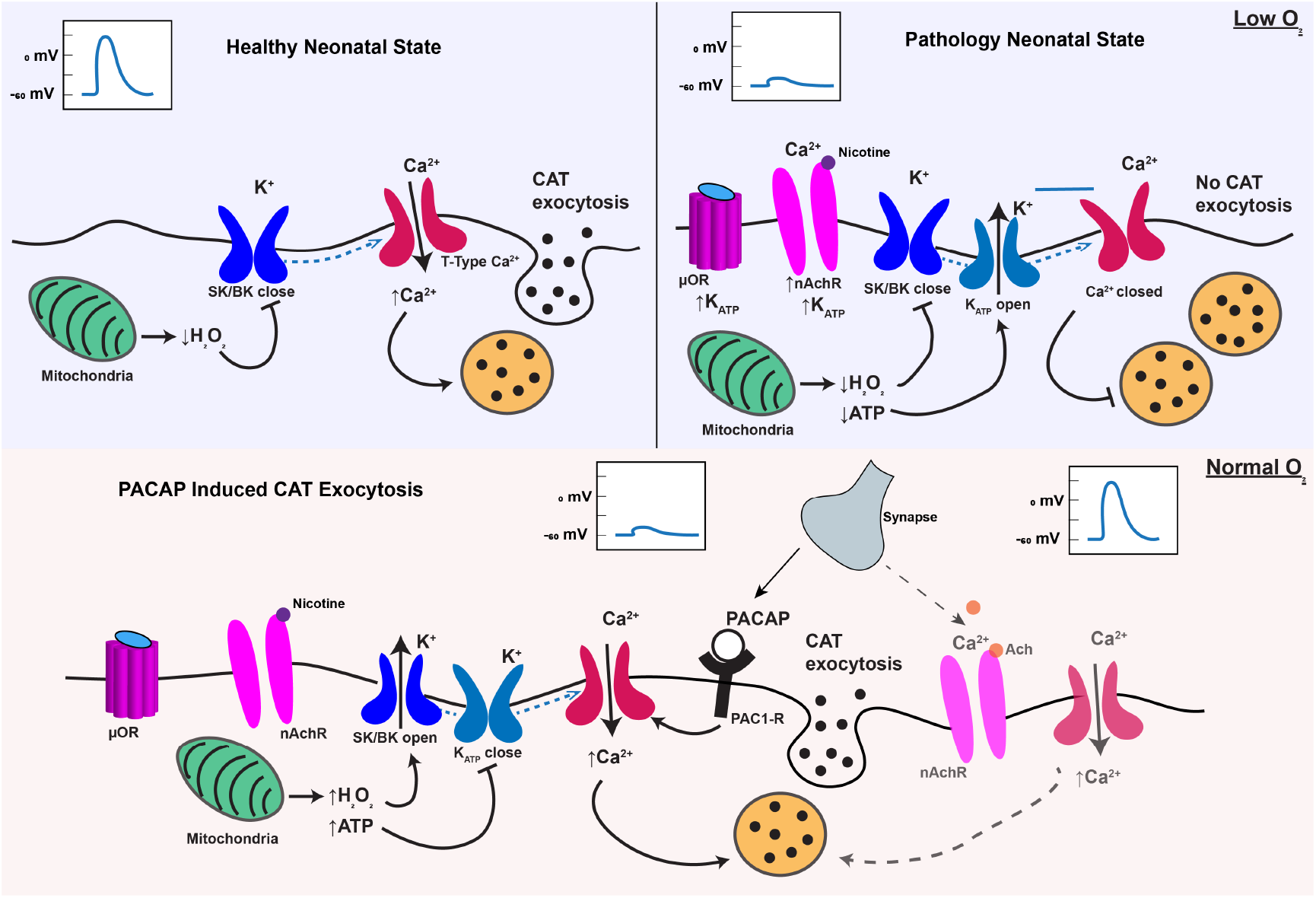
Schematic representation of potential molecular pathways mediating CAT exocytosis in response to normoxia and hypoxia in healthy and drug-exposed neonatal AMCCs.

Despite several indications that adrenocortical outputs are mediated in response to addictive substances in adults, the role of the SAM axis has received little attention (59). Although CATs do not cross the blood-brain barrier, peripheral EP from the adrenal medulla can indirectly alter brain function and behavior by activating vagal afferents (60). While physiological concentrations range from 0 to 0.7 nM for EP and 0.4 to 10 nM for NEP, the ratio of secreted EP/NEP shifts throughout life, resulting in higher concentrations of EP (61). Therefore, to recapitulate adrenomedullary development on-chip, DEX concentrations may be increased over time (62). Future efforts will incorporate additional adrenal cortical cell populations to better mimic the native physiology of the adrenal medulla. Thus, the development of a MPS that recapitulates neuroeffector control systems may be used to better understand SAM regulation in response to acute and chronic drug abuse in adults. Our ‘cut & assemble’ method for rapidly designing and implementing these MPS may provide a clear pathway to elucidate these complex cellular interactions between the nervous system, the adrenal gland, aging, and the ramifications of chronic drug use.

The compartmentalized microphysiological devices developed for adrenal innervation herein may be used to investigate other innervated tissue systems. The nervous system plays a critical role in relaying control and sensory signals from all tissues in the body, and therefore needs to be considered when developing MPS of human organs. The use of GelPins to control the spacial location of cell-laden hydrogels represents a significant development in the organ-chip field as an approach to model anisotropic tissue architectures and innervation (37). Further, these devices allow port threading to add user-friendly fluidic valves and connectors (i.e., Luer Fittings), facilitating the addition of serial or parallel organ systems to establish ‘circuits-on-a-chip’ (36). These fluidic connectors further enable repetitive solution exchanges, while the device’s size reduces CAT dilutions to more easily detectable levels and minimizes the use of cells and reagents required. For example, the impact of sympathoadrenal hormones on CM beating, vascular smooth muscle contraction, and skeletal muscle function may be investigated on the benchtop. In concert with recent developments in human stem cell and organoid culture, these platforms hold the potential to disrupt translational medicine by providing reductionist and physiomimetic test beds.

## MATERIALS AND METHODS

### Device design and fabrication

Custom microfluidic chips were fabricated using a ‘cut and assembly’ method (36) via a commercial laser engraver system (Epilog Zing 16, Epilog Laser) to cut and shape double-sided adhesive tape (966 adhesive, 3M), sheets of polymethyl methacrylate (PMMA, McMaster-Carr) and polyethylene terephthalate (PET, McMaster-Carr), and a polycarbonate (PC) track-etched membrane (Whatman^®^ CycloporeTM, GE Healthcare). These devices use PET GelPins to compartmentalize cell-laden 3D hydrogels in the basal channel and PC membrane to enable culture media diffusion from the apical channel and mimic circulation. Each layer of the chip was designed using computer-aided drafting (CAD, Autodesk Invertor) to have four alignment holes to facilitate layer-by-layer assembly. CAD drawings were exported as vector art into Illustrator (Adobe) and then printed on a laser engraver using material-specific speed and power settings to cut each chip layer. During laser cutting, the protective paper lining on both the PMMA sheets and the 3M tape was left to minimize exposing the materials to burning products. From top to bottom, each of the layers was cut and prepared as described below.

The 4.5 mm PMMA top layer featured two additional 4 mm diameter through-holes, which serve as fluidic inlets and outlets for the apical flow channel, four 1 mm diameter through-holes that enable gel loading into the basal channel, and four circular through-holes present in each layer for alignment. Once cut, the fluidic inlet and outlet ports were hand tapped (10-32 UNF), the protective paper liner removed, and the acrylic cleaned using detergent (Contrad 70, Fisher) and 100% isopropyl alcohol (Fisher). The second layer, or the apical flow channel, was comprised of a 1.5 mm PMMA sheet that was sandwiched between ~60 μm thick double-sided adhesive tape and features a 15 × 15 mm square cutout connected to circular inlets and outlet holes that match those holes in the uppermost PMMA layer, in addition to the four alignment and four gel loading holes. The third layer, or the PC membrane with 1 μm diameter pores, was cut from a 47-inch diameter disk into a rectangle with the same dimensions as the chips and contained the four alignment holes and four gel loading holes to enable access to the basal gel compartment. Together, the four, fifth, and sixth layers comprise the basal gel compartment that contains four discrete areas within a contiguous channel. In addition to the four alignment holes, layers four and six, prepared from one 3M tape (~60 μm) and two sandwiched 3M tapes (~120 μm), respectively, each contains a 15 × 15 mm square cutout connected to each gel filling port with a 1 mm channel. Layer five, or the GelPin layer, was cut from a ~75 μm PET film and featured the same cutouts as layers four and six, but with two interconnected concentric rings of material with radiuses of 5 and 10 mm and a width of ~200 μm remaining in the center of the 15 × 15 square. The seventh and final layer was a 1.5 mm PMMA sheet with four alignment holes or a No. 1 glass coverslip for improved imaging resolution.

The cut layers were assembled layer-by-layer using a custom jig containing alignment posts to facilitate feature alignment. The double-sided adhesive between layers served to simultaneously bond each of the layers together during this assembly process. Once fully assembled, chips were stored under vacuum at 50 °C overnight to eliminate outgassing induced bubble formation during cell culture. Finally, devices were UV sterilized (Spectrolinker XL-1000, Spectronics Corporation) for 600 s on each side before cell culture use.

### Primary adrenomedullary chromaffin cell isolation

Primary AMCCs were isolated from the adrenal glands of two-day-old (p2) Sprague-Dawley neonatal rats (Charles River). Rat pups were euthanized by decapitation per Northeastern’s Institutional Animal Care and Use Committee (IACUC) guidelines. While tissue from ~10 rat pups was collected, adrenal glands were harvested and kept in Hibernate^®^-A (BrainBits, LLC) on ice to ensure cell viability. The fibrous capsule around the isolated adrenal glands was carefully removed so that only the adrenal cortex and medulla remained. Due to each gland’s small size and the indiscrete interface between the cortex and medulla, the whole gland is used to increase tissue yield. Dissected adrenal glands were then placed in a 2 mg/mL collagenase I (~250 units/mg, Gibco) solution containing 0.1% (v/v) trypsin (Gibco) in Hank’s balanced salt solution (HBSS, Gibco) at 37 °C to dissolve the connective tissue and release the AMCCs. After 45 min, the enzymatic digestion was halted by dilution with HBSS, and the tissue further dissociated mechanically via trituration with a fire-polished glass pipette. A 70-μm cell strainer was then used to remove any remaining undigested tissue from the solution. The dissociated cell suspension was centrifuged at 300 g for 5 min, resuspended in complete culture media [Dulbecco’s Modified Eagle Medium (DMEM, Sigma) supplemented with 10% fetal bovine serum (FBS, Sigma), 2 mM L-glutamine (Gibco), and 50 units/mL penicillin/ streptomycin (Gibco)], and then counted for use within one hour following the dissociation. To confirm successful isolation and dissociation, AMCCs were seeded on laminin-coated coverslips (~70 μL at 50 μg/mL laminin) and cultured for five days before being stained for chromogranin A (CHGA) and phenylethanolamine N-methyltransferase (PNMT).

### Primary preganglionic sympathetic neuron isolation

Primary p2 neonatal rat PSNs were isolated from the sympathetic division of the spinal cord (T8-L1). To identify the spinal cord’s sympathetic region, the entire spine was first trimmed away from the carcass with portions of the rib cage left attached. Using the ribs as a guide, the cord was severed between the L1 and T8 vertebrae. The column was then opened, and the T8-L1 region of the cord was carefully removed by trimming away attached dorsal root ganglia and other neural projections. While the preganglionic sympathetic neurons are primarily located within the lateral horn, the whole cord was dissociated to increase tissue yield. Further, to improve cell viability, dissected spinal cords were stored in Hibernate^®^-A on ice while the other cords were extracted from ~10 rat pups. The collected spinal cords were minced into ~2 mm pieces and transferred to a 0.2% (v/v) trypsin solution in Hibernate^®^-A without Ca^2+^ (BrainBits, LLC) at 37 °C to dissolve the neural tissue. After 30 min, the tissue suspension was triturated ten times with a fire-polished pipette to break up the tissue further and release the PSNs. The spinal cord digestion was filtered using a 70-μm cell strainer to remove any remaining tissue aggregates.

Smaller myelin debris was removed via density gradient centrifugation (63). In brief, a gradient was prepared such that a 35% (v/v) 1:1 4-morpholinepropanesulfonic acid (MOPS):Opti-prep (Sigma) solution in HBSS was layered on top of a 20% (v/v) 1:1 MOPS:Opti-prep solution in HBSS. The solution of dissociated PSNs was carefully added to the gradient and centrifuged at 725 g for 15 min to separate neurons from myelin debris. Once cellular debris was removed, the PSNs were pelleted (300 g for 5 min), resuspended in complete culture media supplemented with 0.25 μg/mL neural growth factor (NGF, R&D Systems) and 0.1 μg/mL Ciliary Neurotrophic Factor (CNTF, R&D Systems), and then counted for use within one hour following the dissociation. To confirm successful isolation and dissociation, PSNs were seeded on laminin-coated coverslips and cultured for five days before being stained for beta III tubulin, choline acetyltransferase (ChAT), and calretinin. Lastly, a co-culture of AMCCs and PSNs, seed on laminin-coated coverslips, was stained for Synapsin II to demonstrate synaptogenesis between these cell populations.

### Primary cardiomyocyte isolation

Primary CMs were isolated from the ventricles of p2 neonatal rat pups using established protocols (43). In brief, the ventricles were removed and cleaned of all prominent veins, then stored in 0.5% (v/v) trypsin in HBSS overnight at 4 °C. Approximately 20 – 24 hrs later, the cardiac connective tissue was dissolved by sequential collagenase II (305 units/mg in HBSS, Gibco) digestions at 37 °C. A 70-μm cell strainer was used to remove undigested tissue, and CM were enriched via differential attachment. After one hour in a tissue culture flask cultured in standard culture conditions (37 °C, 5% CO_2_) with complete culture media, any unattached cells were considered CMs. Enriched CMs were collected, counted, and seeded at 1000 cells/mm^2^ in tissue culture-treated well plates for later beat rate quantification. CMs were cultured using complete culture media in standard culture conditions with media replaced every other day.

### rO_2_ culture and characterization

As outlined in Fig. S3, equal parts AMCCs and PSNs (1:1) were co-cultured at a total density of 1000 cells/mm^2^ each and compared to mono-cultures of AMCCs and PSNs at 500 cells/mm^2^ and AMCCs at 1000 cells/mm^2^ to investigate the role of innervation in rO_2_ induced catecholamine exocytosis. In a similar context, the role of opioids and nicotine was explored in mono-cultures of AMCCs at 500 cells/mm^2^. The day after seeding, cells were cultured in complete culture media supplemented with 10-5 M arabinoside cytosine (ARA-C, Sigma) for 48 h to remove highly mitotic fibroblasts. Complete culture media was then exchanged every other day. Complete media was supplemented with 1 *μ*M DALDA (Bachem), a mu-opioid agonist, or 1 *μ*M nicotine (Sigma) each time it was replaced to investigate the role of addition drugs. Cultures were left to grow for five days post-seeding under standard culture conditions to allow for synaptogenesis between PSNs and AMCCs or long-term exposure to nicotine and DALDA. AMCC viability was determined via a LIVE/ DEAD^®^ viability/cytotoxicity kit (Life Technologies) after five days in culture.

AMCC hypoxia-induced catecholamine exocytosis was assessed by monitoring changes in CM beat rate. To improve reproducibility, culture media was exchanged 30 min before a 4 hr incubation under either hypoxic condition (37 °C, 7 vol% O_2_) or standard culture conditions. PACAP (100 nM for 30 min, Tocris) was then used as an analog for neurogenic stimuli for understanding AMCC function after exposure to DALDA and nicotine. Changes in CM beat rate caused by the rO_2_-induced or neural-mediated activation of AMCCs was evaluated by transferring AMCC/PSN-conditioned media to the CM cultures and quantifying beat rate after 30 min equilibration at standard culture conditions. Beat rate was quantified with a custom MATLAB code to calculate beating on a cell-by-cell basis using video microscopy (43) as an estimation of CAT exocytosis. Videos were taken from a minimum of three independent cultures with an average of fifteen CMs per video to calculate each condition’s average beat rate. Each experimental group was then done in triplicate using primary cells isolated from separate batches of animals.

### Microphysiological model preparation and culture

To investigate 3D adrenal cell cultures, AMCCs were encapsulated within a GelMA hydrogel. GelMA hydrogel precursor solutions (7.5% (w/v) were prepared in complete culture media with 0.5% (w/v) LAP under sterile conditions and used within 2 h. Dissociated AMCCs were mixed with the hydrogel precursor solution at a density of 4 × 107 cells/mL. Approximately 7 μL AMCC-laden gel precursor solution was photocrosslinked with visible light (405 nm, 10 mW/cm2) between a 180 *μ*L tall spacer and a 3-(trimethoxysilyl) propyl methacrylate (ACROS Organics) coated glass slide for 45 s (0.25 s/μm of hydrogel thickness). AMCC-laden GelMA hydrogels were incubated in standard culture conditions up to 14 days, and complete culture media exchanged every other day. Neurogenetic control of 3D encapsulated AMCC was then assessed as described above.

To prepare an innervated adrenal model, dissociated AMCCs and PSNs were co-cultured in 3D using a photocrosslinkable gelatin-based hydrogel, GelMA, within ccMPSs. Dissociated AMCCs and PSNs were pelleted and resuspended in the GelMA pre-gel solution at a density of 4 × 107 cells/mL and 5 × 106 cells/mL, respectively. GelMA derived from fish gelatin was synthesized as previously reported (43). GelMA pre-gel solutions were prepared at 10% (w/v) in culture media containing 0.5% (w/v) lithium phenyl-2,4,6-trimethyl-benzoyl phosphinate (LAP, Biobots). The AMCC-laden gel precursor solution was carefully loaded into the central chamber of the ccMPS via corresponding the hydrogel fill port so that a liquid-air interface was formed at the GelPin boundary. The AMCC-laden solution was then photocrosslinked in situ with visible light (405 nm, 10 mW/cm2) for 60 s (0.25 s/μm of hydrogel thickness). Following the AMCC-laden hydrogel gelation, the PSN-laden solution was loaded into the microfluidic device and similarly photopolymerized adjacent to the adrenal *μ*Tissue to form a contiguous hydrogel construct with discrete cellular compartmentalization. Complete culture media was perfused into the organ-chips via the fluidic inlet and exchanged daily over 14 days to allow for adrenal innervation.

### AMCC electrophysiology

Dissociated AMCCs were seeded at 1000 cells/mm^2^ on laminin-coated multi-electrode arrays (MEA; Multichannel Systems, 60-6wellMEA200/30iR-Ti-rcr) and cultured for five days as described above. To investigate AMCC electrophysiology, culture media was replaced 30 min before recordings. Extracellular potentials were recorded at 40kHz (MCRack) for 60 – 90 s to establish a baseline. 100 mM of PACAP was added to each MEA, and AMCC electrophysiology was recorded. Culture media was replaced before a final recording. Exocytosis events were sorted and analyzed using an automated threshold sorting feature and visually inspected for artifacts.

### Enzyme-linked immunosorbent assay (ELISA)

AMCC CAT concentrations before and after PACAP stimuli were quantified using an EP and NEP ELISA (Biomatik). Cell culture media was collected from a minimum of three AMCC cultures from two independent experiments for each group tested: AMCC, AMCC + PACAP, AMCC + DEX, AMCC + DEX + PACAP. Collected media was frozen at −80 °C and later quantified on the same EP and NEP plate to converse resources. Differences in light absorption from known EP and NEP concentrations were used to establish a standard curve for interpolating CAT concentrations from AMCC culture media. Increases in CAT concentrations for AMCC cultures with and without DEX in response to PACAP were reported as the ratio of AMCC + PACAP over AMCC for both EP and NEP.

### Immunocytochemistry

Innervated adrenal μTissues and adrenal co/mono-cultures were fixed for 30 min with 4% paraformaldehyde and then permeabilized for 20 min with 0.1% X-100 Triton at room temperature. After permeabilization, samples were rinsed with DPBS and blocked for >12 hrs with 2.5% goat serum in DPBS at 4 °C. Samples were then incubated for an addition 12 – 24 hrs with 1:400 mouse anti-beta III tubulin (Invitrogen, 480011), 1:300 sheep anti-ChAT (Abcam, ab18736), 1:300 mouse anti-nNOS (Santa Cruz, sc5302), 1:300 rabbit calretinin, 1:50 rabbit anti-CHGA (Abcam, ab15160), 1:100 mouse anti-Synapsin IIa (Santa Cruz, sc136086), and 1:100 mouse anti-PNMT (Santa Cruz, sc393995) in newly prepared blocking solution at 4 °C. Following incubation with primary antibodies, samples were washed thrice with DPBS and 5 – 10 min wait steps between rinses. To visualize the primary antibodies, anti-chicken, anti-sheep, anti-mouse, and anti-rabbit fluorescently labeled secondary antibodies were added for an overnight (~12 hrs) incubation at 4 °C. For cell grown on coverslips, samples were again rinsed thrice with DPBS and then mounted on cover slides with ProLong^®^ Gold Antifade with 4’,6-diamidino-2-phenylindole (DAPI). For *μ*Tissues, samples were rinsed ever 3 – 4 hrs for ~12 hrs with DPBS and then incubated with 1:1000 DAPI in DPBS for 15 minutes to stain for cell nuclei before one addition rinsing cycle. All samples were imaged using an inverted fluorescence microscope (Zeiss Axio Observer Z1).

### Statistical analysis

All data were first checked for normality using the Shapiro-Wilk test. Statistical significance was then calculated using GraphPad PRISM 7 using a Tukey one-way ANOVA. All beat rate data were normalized to controls. Error bars represent the mean ± standard deviation of measurements (**p < 0.05).

## SUPPLEMENTARY MATERIAL

Fig. S1

Fig. S2

Fig. S3

Fig. S4

## ACKNOWLEDGMENTS

The authors would like to thank the Department of Chemical Engineering, College of Engineering, Northeastern University for administrative support.

## Funding

American Heart Association (AHA, 19PRE34430181, JR Soucy); National Institute of Health (NIH, R21EB025395-01, RA Koppes), American Heart Association (AHA, 19IPLOI34760604, RA Koppes)

## Author contributions

JRS and RAK conceived the project. JRS synthesized materials, isolated primary cells, fabricated chips, seeded and preformed all cell culture work, stained and imaged all cultures, developed CAD models, analyzed results, collected and processed chromaffin cell electrophysiology data, ran the ELISA, prepared the figures, and wrote the manuscript. GB assisted in adrenal cell cultures. RB assisted in chip fabrication. ANK and DTB provided intellectual input and advice. All authors edited and provided feedback on the manuscript. RAK supervised the work.

## Competing interests

No competing interests to report.

## SUPPLEMENTARY MATERIAL

**Fig. S1.**
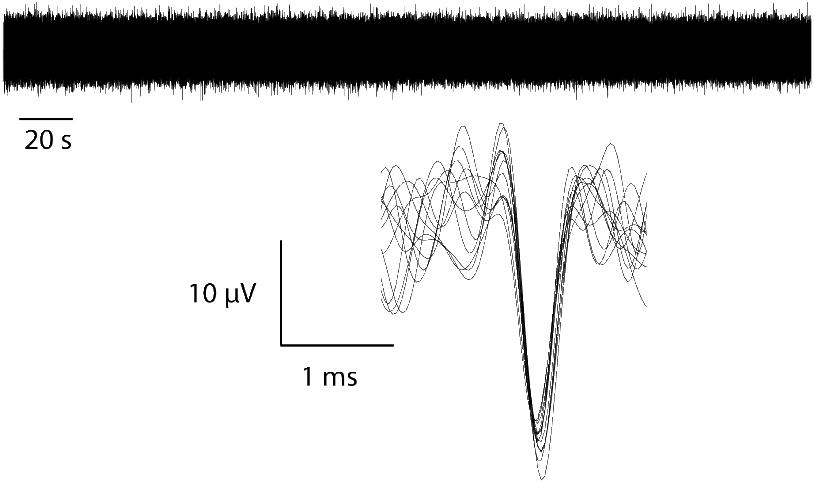
Representative AMCC depolarization.

**Fig. S2.**
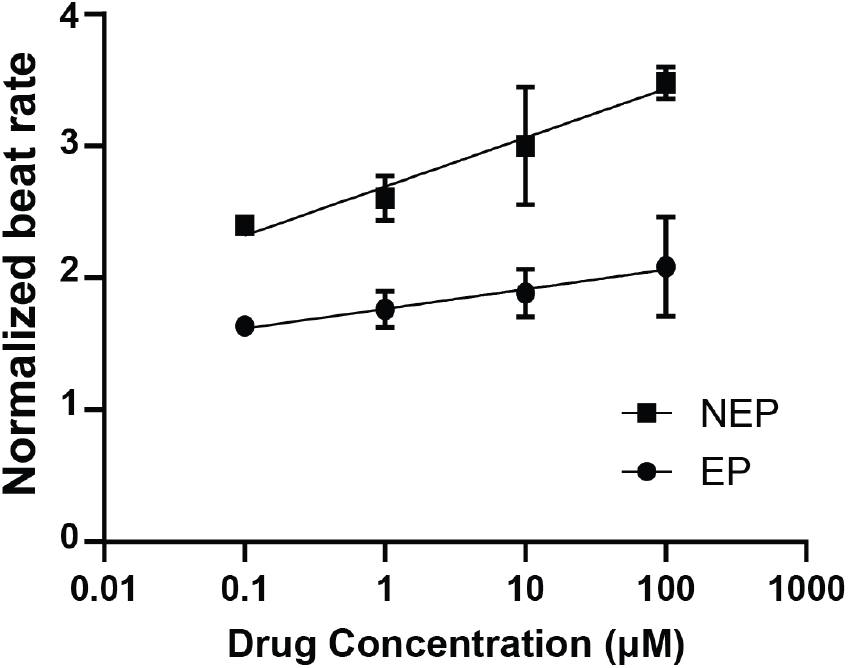
Cardiac dose-response effect. Beat rate increases logarithmically with increasing concentration of CATs.

**Fig. S3.**
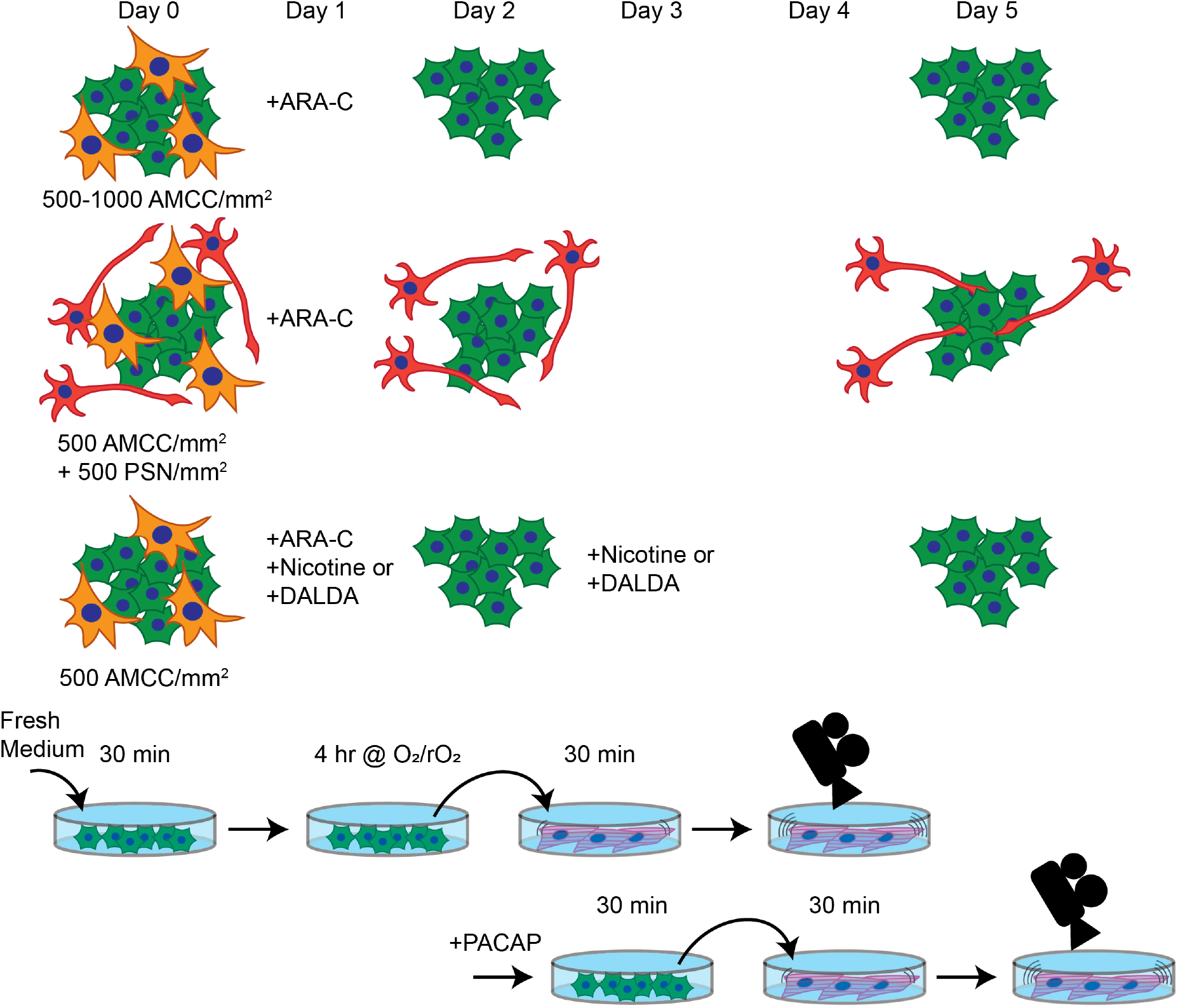
Overview schematic of AMCC culture and CAT quantification via CM beat rate.

**Fig. S4.**
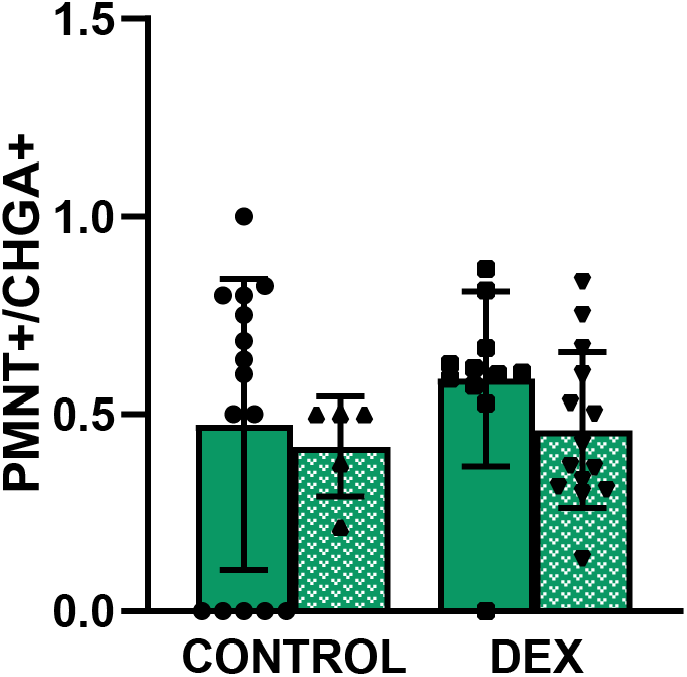
DEX treatment leads to trending increases in the percentage of PNMT positive AMCC populations.

## Notes

### Competing Interest Statement

The authors have declared no competing interest.

